# Ontogeny of Projections from the Motor Cortex Governing Whisker and Neck Movements to the Striatum in the Rat

**DOI:** 10.1101/2023.09.15.558003

**Authors:** Albert T Gu, Victor Z Han, Yanfen Jiang

**Affiliations:** Computational and Systems Biology Program, University of California, Los Angeles, Los Angeles, CA 90095; Center for Integrative Brain Research, Seattle Children’s Research Institute, Seattle, WA 98109 and Department of Biology, University of Washington, Seattle, WA 98195; Division of Gastroenterology, Hepatology, and Nutrition, Department of Pediatrics, Stanford University, Palo Alto, CA 94304

**Keywords:** circuitry, development, postnatal, anterograde tracing, synaptogenesis

## Abstract

Corticostrial or cortico-basal ganglia circuitry plays an important role in integrating sensory and motor information, developing appropriate goal-directed behavior, promoting the maturation of GABAergic interneurons in the striatum, and regulating the nigrostriatal pathways. Dysfunction of this circuitry has been seen in some movement disorders. However, the dynamic changes of this circuitry in early life are not fully elucidated. Previous studies demonstrated that projections from motor cortices of caudal forelimb and jaw-lip-tongue areas to the striatum developed postnatally with terminal-like fibers evident at postnatal day 7. Here we report the development of the projections from the motor cortex governing whisker and neck movements to the striatum. Corticostriatal projections from this area were mainly ipsilateral and also underwent a progressive, postnatal development. The pyramidal tract and its collaterals to the dorsal striatum appeared on the day of birth (postanatal day 0 (P0)), peaked on P6 in density, and continued to be tuned until P36. The intertelencephalic projections in the dorsolateral striatum were established between P6 and P12 and continued to be refined between P20 and P36. Neurons in this motor cortex sent their axons to the contralateral motor cortex via the corpus callosum at the age between P6 and P12. Our results suggest that the time window between P6 and P12 is critical for the development of the projections from the motor cortex governing whisker and neck movement to the striatum in the rat. The overall process of the development of this circuitry appears to correspond to the functional development of whisker movement and locomotor activities.

## Introduction

Movement disorders have a high prevalence. For example, there are about 90,000 people in the US who are diagnosed with Parkinson’s disease (PD) each year, and it is projected that there will be 1.2 million people with PD in the US alone by 2030 (Parkinson’s Foundation, [Prevalence & Incidence | Parkinson’s Foundation](https://www.parkinson.org/understanding-parkinsons/statistics/prevalence-incidence)). In North America, estimates of age-sex-adjusted incidence of PD ranged from 47 to 77 per 100,000 among persons aged 45 and older and increased with age, ranging from 108 to 212 per 100,000 among persons aged 65 and older (Willis AW et al., 2022). Tics, manifested as involuntary, repetitive and sudden twitches, movements, or sounds, are common in young people. Transient tics affect as many as 20% of school-age (5 to 18 years old) children. It was estimated that 4 to 8 cases per 1000 school-age children have Tourette syndrome (Scahill L et al., 2014). Globally, from the limited but increasing data available from regions in low- and middle-income countries, birth prevalence for pre-/perinatal cerebral palsy (CP) was as high as 3.4 per 1000 live births. Birth prevalence for pre-/perinatal CP in regions from high-income countries was 1.5 per 1000 live births, and 1.6 per 1000 live births when postneonatal CP was included (McIntyre S et al., 2022). In the United States, about 764,000 children and adults currently have cerebral palsy, among whom about 500,000 children under age of 18 currently have CP (Prevalence of Cerebral Palsy | Incidence | CerebralPalsy.orgCerebralPalsy.org). Caring for patients with motor disorders each year is a heavy burden for society. For Parkinson’s disease alone, direct healthcare-related, indirect disability expenses and lost productivity in the United States amount to 51.9 billion dollars annually, and the projected total economic burden will surpass $79 billion by 2037 (Yang W et al., 2020). Unfortunately, knowledge about these diseases is limited and current available therapies are not satisfactory. There is a clear unmet medical need for novel therapies.

Disfunction of cortico-basal ganglia circuitry has been observed in multiple movement disorders including Parkinson’s disease, Huntington’s disease, attention-deficit hyperactivity disorder (ADHD), and Tourette syndrome (Crittenden JR and Graybiel AM, 2011), (Shepherd GM, 2013), (Kuo HY and Liu FC, 2019). It is well known that the motor cortex in the frontal cortical area is involved in planning and executing movement behavior via direct corticospinal projections. However, most of the cortical regions also send their projections to the basal ganglia which in turn send outputs to the downstream effector brain regions that are involved in the generation of behavior such as thalamic nuclei, midbrain regions, the pedunculotegmental nucleus, and the hypothalamus.

These cortico-basal ganglia circuits serve as hub for the integration of sensory and motor information and play a central role in developing appropriate goal-directed behavior by transforming cerebral cortex activities into directed behavior such as motor learning, habit formation, and selection of actions based on desirable outcomes (Dudman and Gerfen 2015), (Haber SN, 2016). Developmentally, innervation by the glutamatergic corticostriatal terminals also promotes the maturation of the GABAergic interneurons in the striatum (Sadikot AF and Sasseville R, 1997) and regulates the nigrostriatal pathway (Plenz D and Kitai ST, 1998).

Postnatal development of the corticostriatal projections in the rat has been investigated with anterograde and retrograde tracers including horseradish peroxidase conjugated with wheat germ agglutinin (WGA-HRP) and biocytin. WGA-HRP retrogradely-labeled neurons were found on postnatal day 7 on both the ipsi- and contralateral cortices and these labeled neurons were increased in number and distributed through layers III to VI of the isocortex by day 14 (Iniguez C et al., 1990). After being injected into the caudal forelimb motor cortex, anterogradely biocytin-labeled fibers were found as early as postnatal day 1 in the dorsolateral striatum, denser on the ipsilateral side. These fibers displayed a patch/matrix-like morphology from postnatal day 14 and onward. Similarly, biocytin-labeled fibers were found as early as postnatal day 0, with an increase of terminal-like profiles up to postnatal day 7 in the ventrolateral striatum after biocytin injection in the jaw-lip-tongue area of the motor cortex. Biocytin-labeled fibers from the cingulate cortex appeared to be more developed at birth than the projection from the motor cortex and increased its fiber density in the ipsilateral dorsomedial striatum up to at least postnatal day 7 (Christensen J et al., 1999).

Whisker-based tactile sensation plays an essential role in rodent’s daily life such as navigation, object recognition and social interaction (Sofroniew NJ and Svoboda K, 2015). Diverse innate and learned whisker movements allow rodents to gather specific types of information and trigger specific goal-directed behavior (Esmaeili V et al., 2020). Whisker movements in the rat are known to be governed by medial region of the primary motor cortex which also appear to control neck movement based on an intracortical microstimulation study (Tandon S et al., 2008). In the current study, the postnatal development of the corticostriatal connections from the motor cortex controlling whisker and neck movements was investigated with an in vitro anterograde DiI (1,1′-dioctadecyl-3,3,3′,3′-tetramethylindocarbocyanine perchlorate) tracing combined with confocal microscopy for observation and analysis.

## Materials and Methods

The protocol performed in the present rodent study was approved by the Institutional Animal Use and Care Committee of the Seatle Children’s Research Institute, in accordance with the guidelines of the National Institutes of Health and the US Department of Agriculture for the care and use of laboratory animals. Neonatal male and female Sprague Dawley rats were anesthetized with isoflurane vapor in a Bell jar and perfused on postnatal day 1 (P0), day 3 (P2), day 7 (P6), day 13 (P12), day 21 (P20), and day 37 (P36) with a 4% paraformaldehyde solution, pH 7.4. The brains were immediately removed and stored in the same fixative at 4°C until further processing.

Each brain was blocked and sectioned from rostral to caudal to expose the primary motor cortical area. Each brain block was stained with methylene blue to enhance the visualization of the morphological features of the exposed surface of the block, as described elsewhere (Hutton LA et al., 1998). A short acupuncture needle was used to place a small crystal of DiI (Molecular Probes, Eugene, OR, a generous gift from Dr. Hongwei Dong, University of Southern California, Los Angeles, CA) into the layer V of the rostral primary motor cortex in the left or right hemisphere at the level equivalent to AP 1.2-1.45 in the adult brain (Swanson, 2004) under visual guidance with the assistance of a dissecting microscope. For all implants, DiI crystals selected were similar in size as judged by eye when viewed under a dissecting microscope. The implanted brain blocks were stored in 0.4% paraformaldehyde, and DiI was allowed to diffuse for 2.5 months in the dark at 37°C. The brain blocks were then embedded in 1.5% agarose, and 100-μm-thick sections were cut on a Leica vibratome, mounted onto poly-

L-lysine-coated glass slides, and coverslipped with 0.01M phosphate buffer, pH 7.4. Sections were evaluated with both conventional fluorescence and confocal microscopy. Confocal images of DiI-labeled fibers were collected through the ipsi- and contralateral striatum and the contralateral motor cortex by using a Zeiss LSM-710 confocal microscope operated by Zen software. For quantification, a series of 15 adjacent optical sections (20× objective, 1.69 um-Z interval) were projected and image analysis was performed by using ImagePro Premier software (Media Cybernetics, Rockville, MD). The total area of pixels in each image was counted and used as an index of the overall density of labeled fibers in the observed optical field.

## Results

Brains with DiI crystals implanted in the primary motor cortex governing whisker and neck movements (Figure 1 A) were obtained from P0 (n=4), P2 (n=3), P6 (n=5), P12 (n=5), P20 (n=3), P36 (n=6) male and female rats. The crystalline deposits of DiI were aimed to be confined in layer V (Figure 1A). Shortly after implantation, DiI crystals started to dissolve, and diffusion of the dye was observed on the dorsum of the brain days after implantation which expanded over time and reached to the adjacent secondary motor area (Figure 1B). Slight differences in diffusion size have been observed, likely due to the slight differences in DiI crystal sizes.

**Figure 1.**
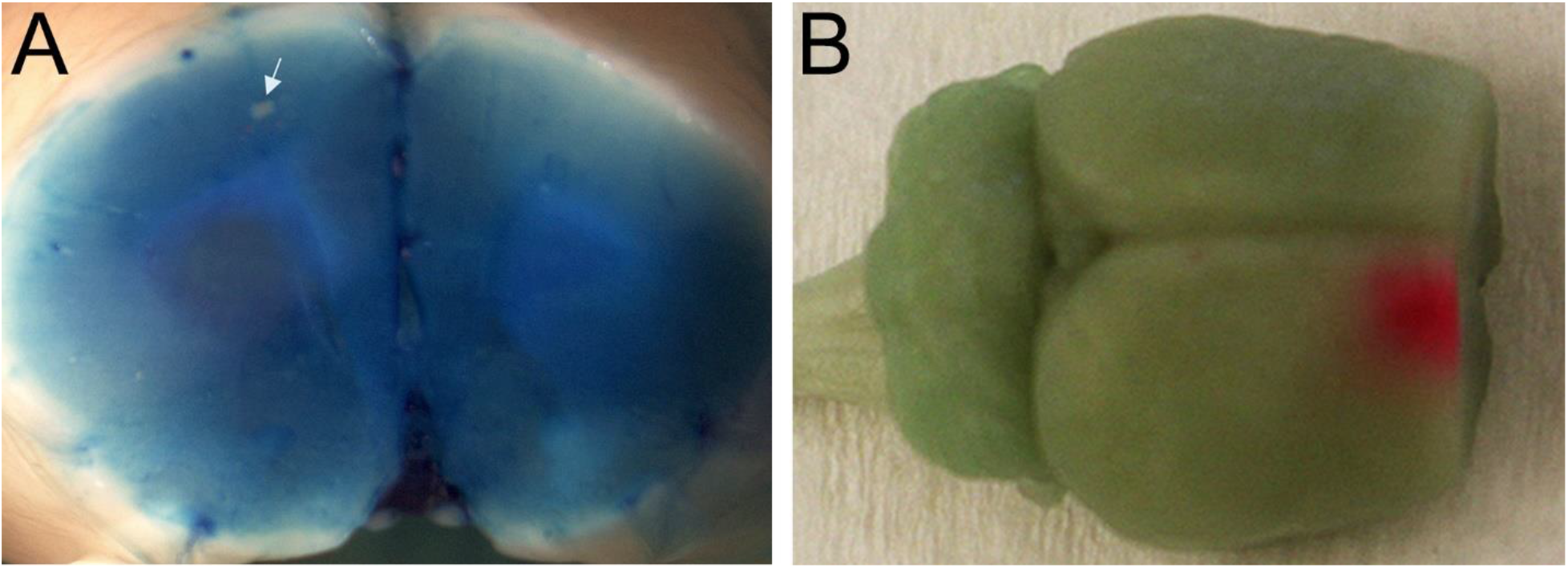
A. Location of DiI implantation (arrow). B. Diffusion of DiI (red) tracer in the motor cortex 10 days after implantation. DiI crystal slowly melted and diffused in the brain tissue.

Cases with the maximum labeled fibers at each postnatal age were selected for analysis. At the most mature age (P36) studied, the corticostriatal projections were predominantly localized to the ipsilateral dorsal and dorsolateral part of the striatum (Figure 2A). Therefore, sampling at the regions of 12 o’clock and 9 or 3 o’clock was used to depict the locations of the corticostriatal projections in the dorsal and dorsolateral regions, respectively, in the current study (Figure 2B).

**Figure 2.**
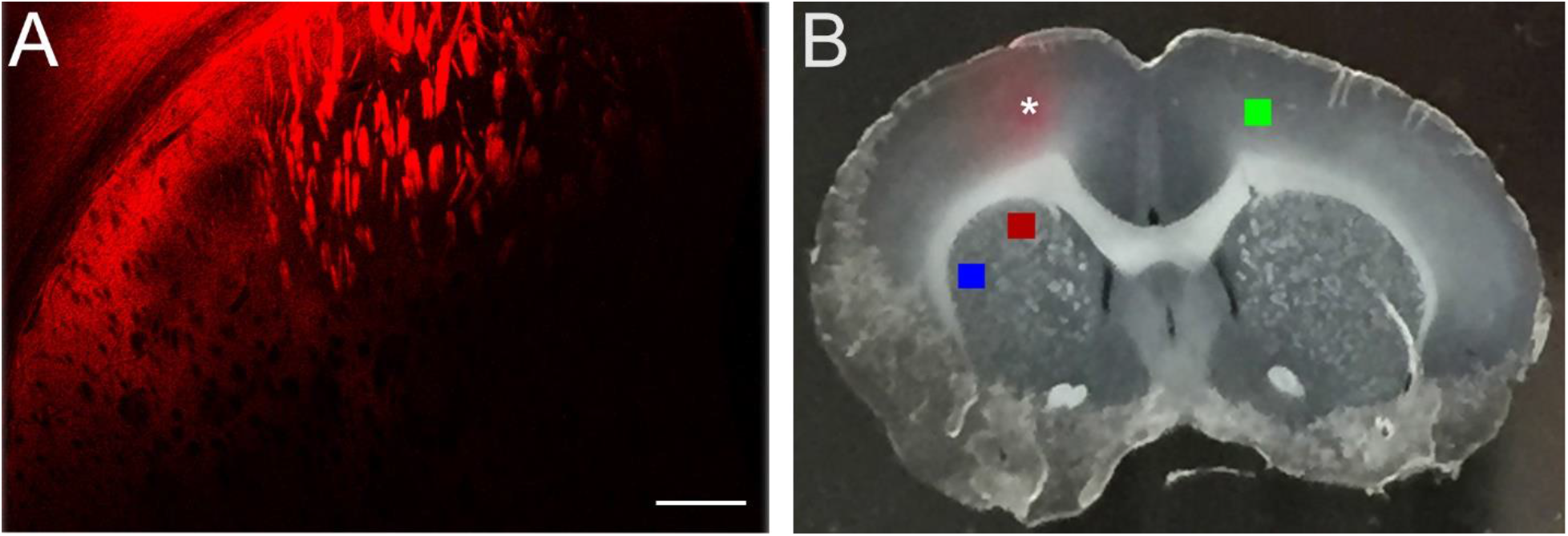
A. Low power confocal microscopic image of a P36 brain indicating that corticostriatal fibers are located in the dorsal and dorsolateral striatum. B. DiI in the motor cortex (asterisk), and the sites of at 12 O’Clock (red rectangle), 9 O’Clock (blue rectangle), and contralateral motor cortex (green rectangle) where confocal images were captured. Scale bar = 200 μm.

Patches of DiI-labeled fibers in the pyramidal tracts started to show at the 12 O’clock location in the striatum on P0. The density of these fibers appeared to increase quickly at the postnatal ages observed, and underwent refinement evidenced by slight reductions in numbers of patches, and the changes in sizes and smoother borders of the pyramidal tracts on P36. Similarly, fine DiI labeled terminals that are supposed to make direct contacts with the dorsal striatal neurons in between the pyramidal patches had the similar development pattern – from sparse density on P0 to higher density on P2 till P12, and to a plateau in density from P20 to P36 (Figure 3A-F, Table 1). DiI-labeled fibers at 9 or 3 o’clock of the ipsilateral striatum were sparse on P0, P2, and P6 (Figure 4A, B.C), but gained a significant increase in density on P12 (Figure 4D). The density of the labeled fibers continued to increase and peaked on P20 (Figure 4E, Table 1). On P36, there was decrease in fiber density and these fibers became finer and more punctate (Figure 4F).

**Table 1.** Pixel areas of corticostriatal fibers at different postnatal ages.

|  | P0 | P2 | P6 | P12 | P20 | P36 |
| --- | --- | --- | --- | --- | --- | --- |
| 12 O'clock,<br>Dorsal striatum | 273147 | 361267 | 494884 | 709477 | 683289 | 719792 |
| 9 or 3 O'clock,<br>Dorsolateral<br>striatum | 3776 | 2880 | 3640 | 27566 | 366312 | 275710 |
| Contralateral<br>motor cortex | 11 | 78 | 11 | 616903 | 672994 | 494704 |

**Figure 3.**
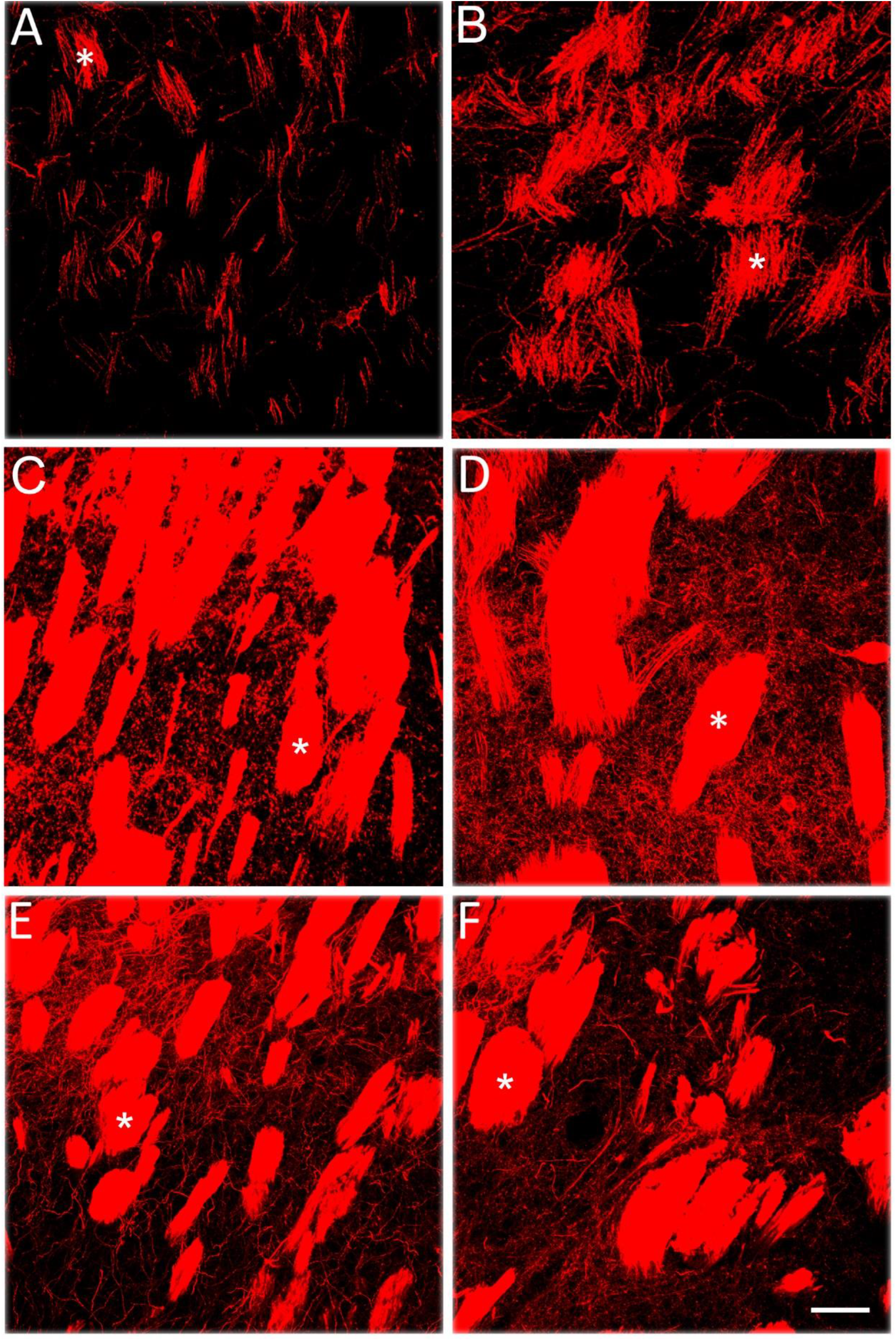
Confocal images of DiI-labeled fibers at the dorsal striatum at P0 (A), P2 (B), P6 (C), P12 (D), P20 (E), and P36 (F). Asterisks indicate pyramidal tracts. The pyramidal tracts appeared on P0 and rapidly grew in bundle sizes. Large amount of fine terminals, emerged at P6. Scale bar = 50 μm.

**Figure 4.**
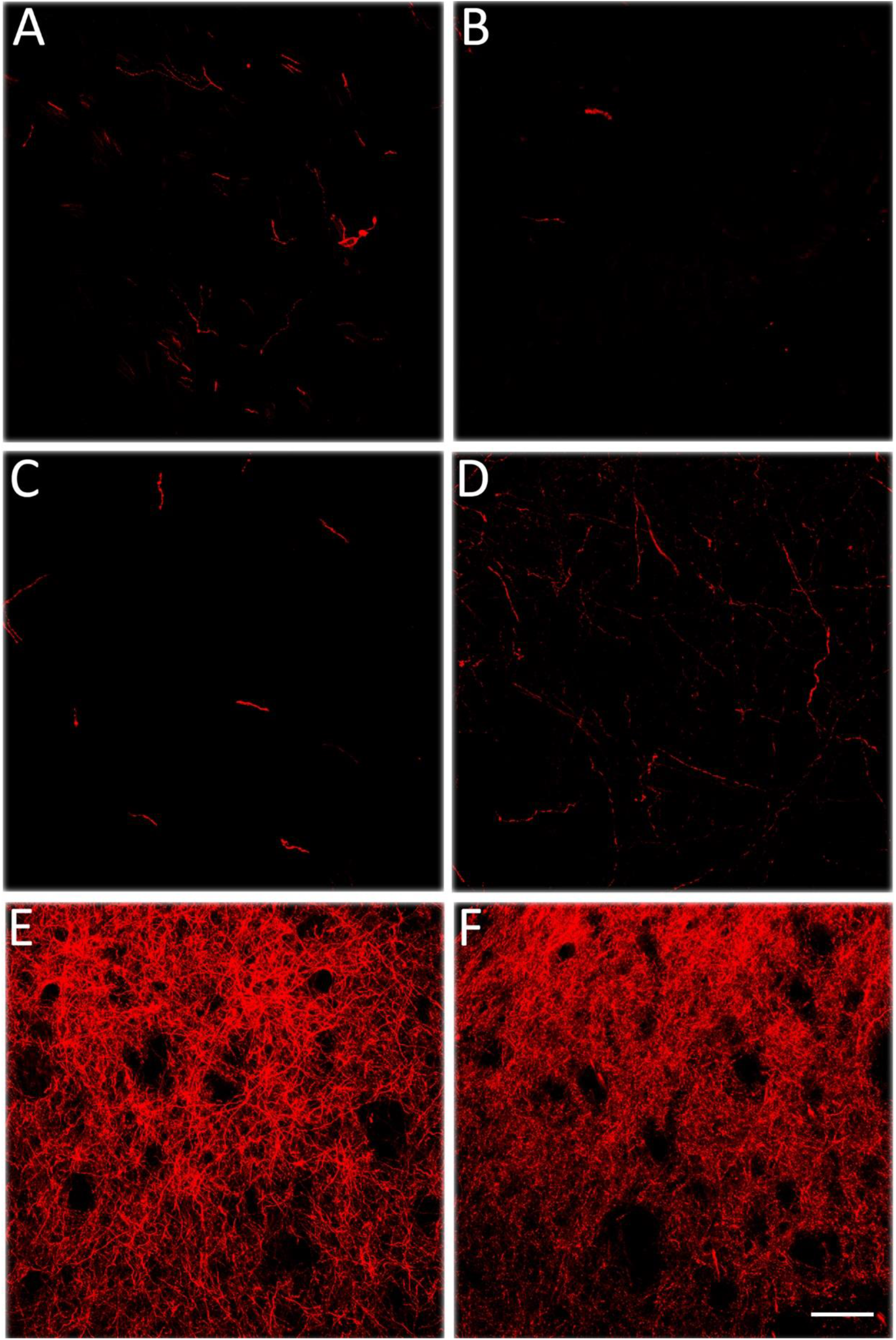
Confocal images of DiI-labeled axons at the dorsolateral striatum at P0 (A), P2 (B), P6 (C), P12 (D), P20 (E), and P36 (F). Good amount of labeled axons appeared at postnatal day 6 and continued to increase in density at older age. Scale bar = 50 μm.

DiI-labeled fibers were found in the corpus callosum as early as P2. Some of them appeared as axonal growth cones in morphology, but no fibers were seen to cross the midline (Figure 5A) at this age. Labeled fibers that crossed midline started to be seen on P6 and increased in density at later postnatal ages (Figure 5B). Crossed fibers made their stop at the contralateral motor cortex on P12. Density of these fibers appeared to increase on P20, and became finer and more punctate with a decrease in overall density on P36, in a way similar to the profiles at 9 or 3 o’clock of the ipsilateral striatum (Figure 6A-F, Table 1). A small number of crossed fibers also ran ventrally and stopped at the contralateral dorsal and dorsolateral striatum after P20 (data not shown).

**Figure 5.**
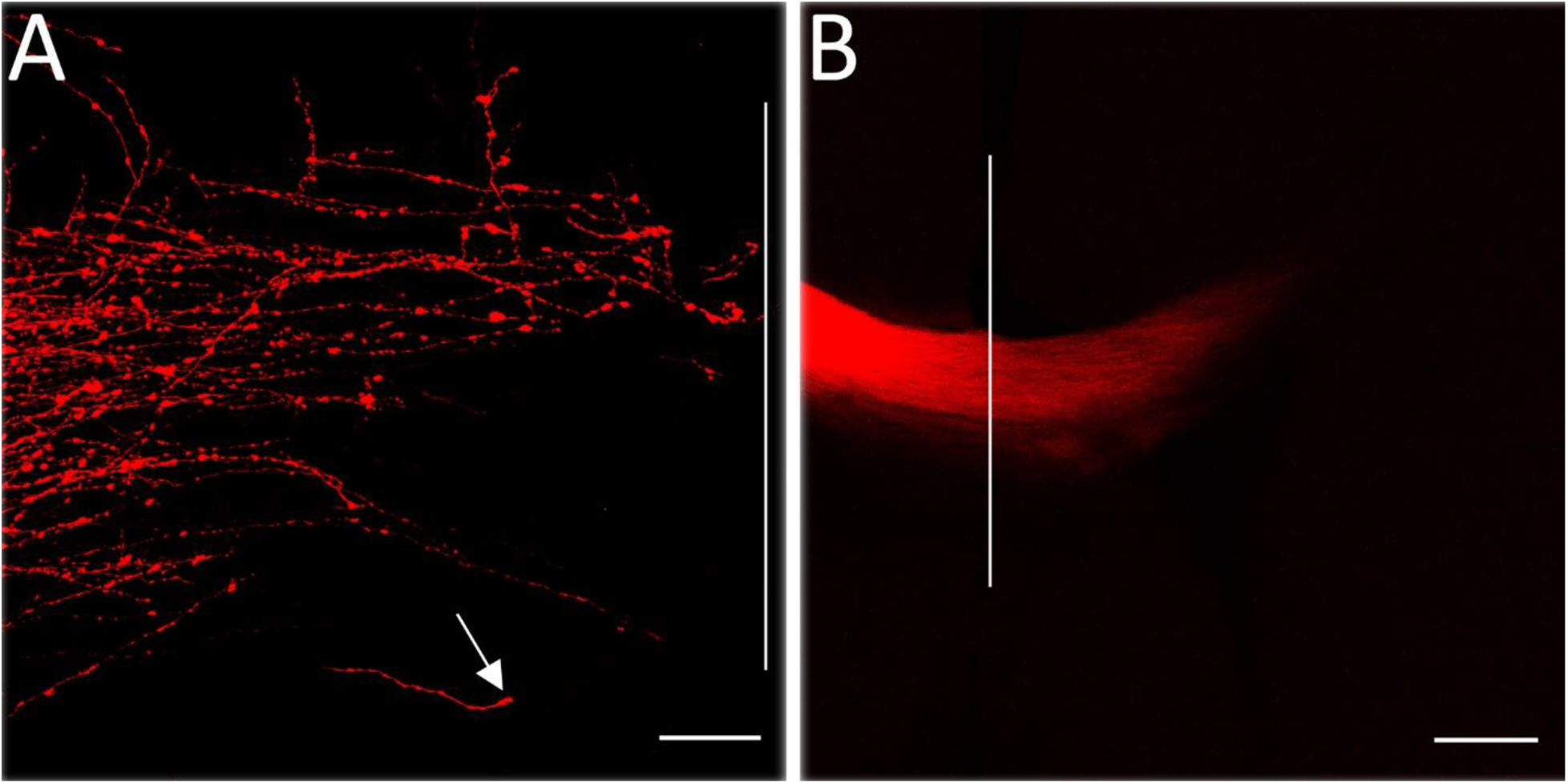
Confocal images of the DiI-labeled axons in the corpus callosum at P2 (A) and P20 (B) brains. Arrow indicates a growth cone of an axon, and vertical lines indicate the midline of the brains. Fibers appeared to cross the midline between the time window of P2 and P6. Scale bar = 15 μm (A) = 200 μm (B).

**Figure 6.**
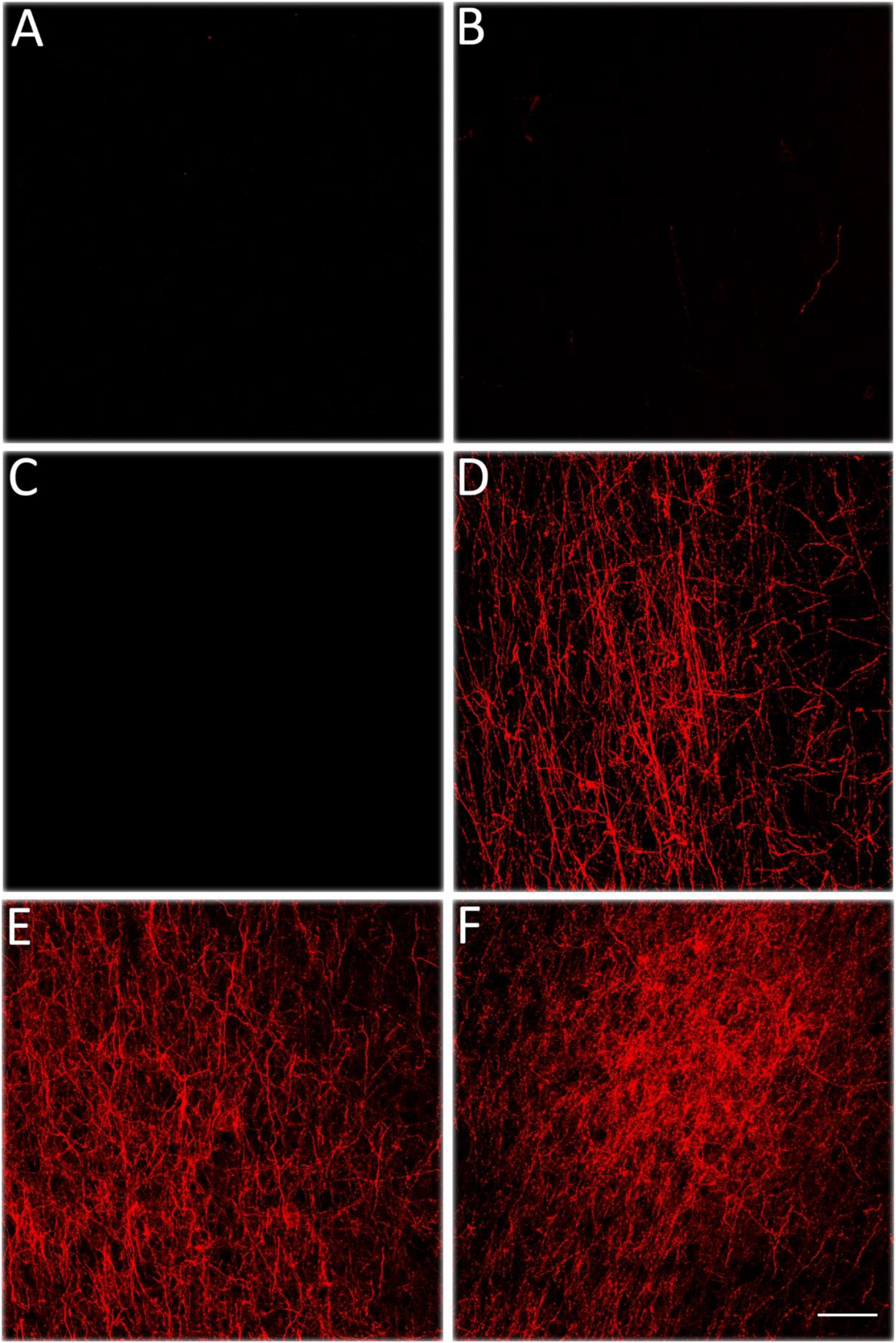
Confocal images of DiI-labeled axons at the contralateral motor cortex at P0 (A), P2 (B), P6 (C),P12 (D), P20 (E), and P36 (F). The motor cortex to contralateral motor cortex projections appeared to develop between P6 to P12 and continued to increase in density at older age. Scale bar = 50 μm.

## Discussion

Consistent with previous results (Iniguez C,De Juan J,al-Majdalawi A and Gayoso MJ, 1990), (Christensen J,Sorensen JC,Ostergaard K and Zimmer J, 1999), the current study indicates that corticostriatal projections from the motor cortex governing whisker and neck movement started to develop early in life, sparse labeled fibers were seen as early as P0 in both dorsal and dorsolateral regions of the striatum, similar to the projections from other regions of the motor cortex. The topographic pattern and the time course revealed in the current study resemble those of the corticostriatal projections from the caudal forelimb area of the motor cortex (Christensen J,Sorensen JC,Ostergaard K and Zimmer J, 1999).

Differences in morphology of labeled fibers and time course have been observed between the dorsal and dorsolateral regions of the striatum. The dorsal region harbors the pyramidal tracts containing corticospinal and cortico-brainstem fibers. Thin pyramidal tracts started to be seen in the dorsal striatum on P0 and gradually increased in size, peaking on P12. Individual tracts then evolved in size and became smaller since P20. Correspondently, labeled terminals arborized from the pyramidal tracts to end in the areas between the pyramidal tracts. These terminal-like profiles became robust since P6 and evolved to be finer and more punctate thereafter. In the dorsolateral striatum, however, except for labeled terminals, there were no labeled pyramidal tracts in this region. The labeled fibers appeared to be a part of the intratelencephalic tracts and only a very small amount of them were seen at P0, P2, P6, and P12. Active development happened late - robust labeled terminals were observed at P20, and the density of these terminals appeared to peak at this age. The terminals became finer afterwards when the pups grew older. The terminals in the dorsal and dorsolateral regions may come from different types of neurons in the motor cortex as suggested in the previous report (Reiner A et al., 2003).

The development pattern of the pyramidal tract and the corticostriatal circuits demonstrated in the current study appears to correspond to the functional development of whisker movement (Grant RA et al., 2012). It was found that from as early as P2 there were some small movements of the whiskers. From P6 onwards, whisker movement became more frequent and retraction movements started to occur, in parallel with increased movement of the head and limbs. Between P11 and P15, bilateral rhythmic whisking emerged, alongside the emergence of walking, followed by the first signs that whisker movements were being modulated by environmental contacts. During this period, pups began to locomote in a forward direction effectively. From P15, sensory regulation of whisker movement with the establishment of adult-like patterns of contact-related asymmetry and asynchrony and alterations in whisker spread was accomplished, to match the maturity of animal’s locomotion capabilities. This would allow the animal to accurately control environmental contacts and significantly boost the information gained through vibrissal touch (Grant RA,Mitchinson B and Prescott TJ, 2012).

The early appearance of labeled pyramidal tracts in the dorsal striatum on P0 suggests a prenatal development of corticospinal and cortico-brainstem pathways in the rat because of the long distance. Previous study demonstrated that corticospinal axons reached upper cervical spinal cord levels at P0, reached third thoracic spinal cord segment at P1, extended into the eighth thoracic segment at P3, into the first or second lumbar segment at P7 and into the second to third sacral segment at P9 (Gribnau AA et al., 1986). On the contrary, based on the results demonstrated in this study, the intratelencephalic corticostriatal projections to the dorsolateral region of the striatum appeared to gain speed of development postnatally. The exact mechanism of this late development is unknown but may be due to the late development of the overall locomotion and goal-directed behavior.

Our findings are also consistent with the previous results (Christensen J,Sorensen JC,Ostergaard K and Zimmer J, 1999) that motor cortex governing whisker and neck movement only sent very small amount of terminals to the dorsolateral region of the contralateral striatum, in contrast to the robust projections to the contralateral motor cortex. The development of the cortico-cortical (contralateral) connections appeared to happen postnatally. It takes at least 2 days for the axons in the corpus callosum to cross the midline and approximately 12 days for the terminals to populate in the contralateral motor cortex. Refinement of these terminals takes weeks afterwards, in concert with the development of locomotor and whisker movements.

Knowing how the motor system in the brain is wired in early life and the associated molecular mechanisms may shed light to the discovery of new targets for therapeutic interventions. In many mammals including humans, cats, and rats, sophisticated motor behavior gradually develops after birth. It takes years for humans to develop motor skills such as walking without assistance, jumping etc., and it takes about a month for rats to develop mature motor skills. Corresponding to the development of these movement functions, refinement of the projected terminals from the motor cortex to the striatum and the contralateral cortex has been observed at the light microscopic level in our study, as well as in other studies (Christensen J,Sorensen JC,Ostergaard K and Zimmer J, 1999), which may reflect the dynamic process of synaptogenesis. The overall process of the development of corticostriatal connections appears largely similar to the development of corticospinal system, in which 3 stages were identified: 1. growth of axons to the spinal cord gray matter during late prenatal or early postnatal period; 2. refinement of the terminations in the gray matter including both elimination of transient terminations and growth of new terminals to nearby targets; 3. motor control development characterized by loss of transient terminations, established growth of axons to local targets, as well as myelination (Martin JH, 2005). Current study suggests that the time window between P6 and P12 is critical for the development of the motor neuron connections with the striatum, as well as the contralateral motor cortex in rats.

Although the physical distance between the motor cortex and the striatum is much shorter than that between the motor cortex and the spinal cord, the time needed to build up both connections is approximately similar.

## Abbreviations

ADHD: attention-deficit and hyperactivity disorder
CP: cerebral palsy
DiI: 1,1′-dioctadecyl-3,3,3′,3′-tetramethylindocarbocyanine perchlorate
GABA: gamma-aminobutyric acid
WGA-HRP: horseradish peroxidase conjugated with wheat germ agglutinin

## Author contributions

AG was involved in study design, experiment, data capture and analysis, and writing of the original draft of the manuscript; VH contributed to brain tissue harvesting, data interpretation, and manuscript reviewing; YJ conceived the study, contributed to the study design and data interpretation, and mentored AG throughout the whole project

## Acknowledgments

We thank Dart NeuroScience, LLC for access to instruments and confocal microscope facility

## Declaration of interests

The authors declare no competing interests.

## Notes

### Competing Interest Statement

The authors have declared no competing interest.

## References

Christensen J, Sorensen JC, Ostergaard K, Zimmer J (1999), Early postnatal development of the rat corticostriatal pathway: an anterograde axonal tracing study using biocytin pellets. Anat Embryol (Berl) 200:73–80.

Crittenden JR, Graybiel AM (2011), Basal Ganglia disorders associated with imbalances in the striatal striosome and matrix compartments. Front Neuroanat 5:59.

Dudman JT, Gerfen CR (2015), The Basal Ganglia. In: Paxino G, editor, Rat Nervous System, 4th ed, Amesterdam: Elsevier. p391–440.

Esmaeili V, Tamura K, Foustoukos G, Oryshchuk A, Crochet S, Petersen CC (2020), Cortical circuits for transforming whisker sensation into goal-directed licking. Curr Opin Neurobiol 65:38–48.

Grant RA, Mitchinson B, Prescott TJ (2012), The development of whisker control in rats in relation to locomotion. Dev Psychobiol 54:151–168.

Gribnau AA, de Kort EJ, Dederen PJ, Nieuwenhuys R (1986), On the development of the pyramidal tract in the rat. II. An anterograde tracer study of the outgrowth of the corticospinal fibers. Anat Embryol (Berl) 175:101–110.

Haber SN (2016), Corticostriatal circuitry. Dialogues Clin Neurosci 18:7–21.

Hutton LA, Gu G, Simerly RB (1998), Development of a sexually dimorphic projection from the bed nuclei of the stria terminalis to the anteroventral periventricular nucleus in the rat. J Neurosci 18:3003–3013.

Iniguez C, De Juan J, al-Majdalawi A, Gayoso MJ (1990), Postnatal development of striatal connections in the rat: a transport study with wheat germ agglutinin-horseradish peroxidase. Brain Res Dev Brain Res 57:43–53.

Kuo HY, Liu FC (2019), Synaptic Wiring of Corticostriatal Circuits in Basal Ganglia: Insights into the Pathogenesis of Neuropsychiatric Disorders. eNeuro 6.

Martin JH (2005), The corticospinal system: from development to motor control. Neuroscientist 11:161–173.

McIntyre S, Goldsmith S, Webb A, Ehlinger V, Hollung SJ, McConnell K, Arnaud C, Smithers-Sheedy H, et al. (2022), Global prevalence of cerebral palsy: A systematic analysis. Dev Med Child Neurol 64:1494–1506.

Plenz D, Kitai ST (1998), Regulation of the nigrostriatal pathway by metabotropic glutamate receptors during development. J Neurosci 18:4133–4144.

Reiner A, Jiao Y, Del Mar N, Laverghetta AV, Lei WL (2003), Differential morphology of pyramidal tract-type and intratelencephalically projecting-type corticostriatal neurons and their intrastriatal terminals in rats. J Comp Neurol 457:420–440.

Prevalence & Incidence, Parkinson’s Foundation, Prevalence & Incidence | Parkinson’s Foundation

Prevalence of Cerebral Palsy, Prevalence of Cerebral Palsy | Incidence | erebralPalsy.orgCerebralPalsy.org

Sadikot AF, Sasseville R (1997), Neurogenesis in the mammalian neostriatum and nucleus accumbens: parvalbumin-immunoreactive GABAergic interneurons. J Comp Neurol 389:193–211.

Scahill L, Specht M, Page C (2014), The Prevalence of Tic Disorders and Clinical Characteristics in Children. J Obsessive Compuls Relat Disord 3:394–400.

Shepherd GM (2013), Corticostriatal connectivity and its role in disease. Nat Rev Neurosci 14:278–291.

Sofroniew NJ, Svoboda K (2015), Whisking. Curr Biol 25:R137–140.

Swanson LW. (2004). Brain maps: structure of the rat brain. Amsterdam: Elsevier.

Tandon S, Kambi N, Jain N (2008), Overlapping representations of the neck and whiskers in the rat motor cortex revealed by mapping at different anaesthetic depths. Eur J Neurosci 27:228–237.

Willis AW, Roberts E, Beck JC, Fiske B, Ross W, Savica R, Van Den Eeden SK, Tanner CM, et al. (2022), Incidence of Parkinson disease in North America. NPJ Parkinsons Dis 8:170.

Yang W, Hamilton JL, Kopil C, Beck JC, Tanner CM, Albin RL, Ray Dorsey E, Dahodwala N, et al. (2020), Current and projected future economic burden of Parkinson’s disease in the U.S. NPJ Parkinsons Dis 6:15.

